# The impact of murine cytomegalovirus (mCMV) route and age at infection upon virus spread, immune responses and the establishment of latency

**DOI:** 10.1101/2021.11.29.469758

**Authors:** Christopher P. Coplen, Mladen Jergovic, Elana L. Terner, Jennifer L. Uhrlaub, Janko Nikolich-Žugich

**Author notes:** To whom correspondence should be addressed at P.O. Box 249221, 1501 N. Campbell Ave. Tucson, AZ 85724, USA.

## Abstract

Cytomegalovirus (CMV) is a ubiquitous human virus, which establishes a characteristic lifetime infection in its host. Murine CMV (mCMV) is a widely-used infection model that has been employed to investigate the nature and extent of CMV’s contribution to inflammatory, immunological, and health disturbances in humans. In an effort to assess the role of route and age in modeling hCMV infection in mice, we have performed a comparative analysis of two common experimental modes of infection (intraperitoneal and intranasal) at two different clinically relevant ages (4 weeks, or prepubescent childhood equivalent, and 12 weeks, or young postpubescent adult). We found that while both routes of infection led to similar early viral loads, differential activation of several parameters of innate immune function were observed. In particular, younger, prepubescent mice exhibited the strongest NK activation in the blood in response to i.p. infection, with this trend holding true in NK cells expressing the mCMV-specific receptor Ly49H. Moreover, i.p. infected animals accumulated a larger amount of anti-mCMV IgG and experienced a greater expansion of both acute and latent phase CD8^+^ T cells. This was especially true for young postpubescent mice, further illustrating a distinction in the bloodborne immune response across not only infection routes, but also ages. These results may be important in the understanding of how a more physiologically applicable model of CMV influences immunity, inflammation, and health over the lifespan.

## INTRODUCTION

Cytomegalovirus (CMV) is one of the most ubiquitous human latent persistent viruses, reaching seroprevalence levels of >90% at advanced ages^1^. In immunocompetent subjects, following brief viremia during primary infection, the virus becomes latent in a small subset of broadly distributed, dominantly mesenchymal cell types (CD34+ hematopoietic progenitors, the monocyte-macrophage lineage including dendritic cells, endothelial cells, and perhaps others)^2–4^. After the establishment of this lifelong latent infection, recurrent subclinical micro-reactivation events can lead to transcription and translation of viral proteins, as well as replication and shedding of the infectious virus. While this virus is well tolerated and compatible with healthy aging in the vast majority of individuals, CMV can be deadly or highly harmful to both immunocompromised individuals and the developing fetus^5^.

Murine CMV (mCMV) has been deployed as a widely-used infection model to investigate causality and mechanisms by which CMV infection may contribute to inflammatory, immunological and overall health disturbances in humans. Indeed, many of the well-documented characteristics of human CMV (hCMV) infection are recapitulated during the course of mCMV infection in mice, including viral latency/persistence, reactivation, T cell memory inflation and others^6–9^. However, the two viruses exhibit only partial homology (~78 open reading frames (ORFs) in the central genome out of ~170 total transcribed ORFs), and many hCMV ORFs do not exist or have no homologues in mice^10^. Importantly, some of the health-outcome associations found between hCMV and chronic diseases in some cohorts (e.g. cardiovascular diseases such as large vessel arteritis, type 2 diabetes, etc.) were not found with mCMV, raising the possibility that virus or other differences may account for some of these discrepancies^11^.

While the genetic differences between both the two viruses and the two mammalian hosts cannot be ignored, other easier-to-control-for discrepancies between hCMV natural infection and mCMV experimental infection exist, including those in the time and route of initial infection. The exact route of entry for natural human infection remains unclear. Primary infection via the oral route has been proposed, however, recent evidence points to nasal infection. Specifically, 1) unlike the pH-sensitive virions of CMV^12^, viruses known to infect gastrointestinally typically have non-enveloped, acid-resistant virions in order to resist acid inactivation, 2) while experimental infection via oral gavage can be achieved in mice, this route is markedly inefficient, and the possibility of respiratory infection via virion regurgitation cannot be excluded, 3) no known entry site has been clearly identified during oral infection, including an absence of infection of gastrointestinal cells in models of mCMV transmission via breast milk^13^, and 4) intranasal (i.n.) inoculation yields robust infection, with olfactory neurons positively identified in several studies as a, if not the, primary entry point in the nasal mucosa during both i.n. inoculation and models of natural transmission^14^.

By contrast, the overwhelming majority of studies in mice have been performed in adult postpubescent animals, infected at 6-12 weeks of age, following systemic administration of mCMV dominantly via intraperitoneal (i.p.) and occasionally via intravenous (i.v.) routes^15–17^, generally with virus doses of 10^5–6^ plaque forming units (pfu) of the virus. Both of these necessitate mechanically breaking the tissue barrier in order to introduce the virus systemically. In humans, the first wave of hCMV infections occurs at early timepoints in life (usually in childhood, before puberty) – through nasopharyngeal / oral route via bodily fluids such as saliva and tears, which does not require a physical barrier breach. These differences between the most popular modes of mouse experimental infection and human natural infection have the potential to result in differences in viral spread, control and subsequent pathogenesis, including potential association with chronic diseases^18^, and therefore require to be studied in depth.

To assess the role of route and age in modeling hCMV infection in mice, here we have performed a comparative analysis of two modes of infection (i.p. and i.n.) at two different ages - 4 weeks (prepubescent childhood equivalent) and 12 weeks (young postpubescent adult). We found that both routes of infection yielded similar viral loads (even when lower initial dose was inoculated i.n.), with some potentially interesting differences in virus spread, such as modestly decreased levels of viral burden across each tissue in 4-week old i.n. infected animals. We further found that some parameters of innate immunity may be more robust in i.p. infection, raising the possibility that different arms of immunity may be engaged as a function of virus route of entry. Overall, these results begin to define relationships between the potentially more physiological route and age at infection, and the key parameters of CMV spread, immune control and establishment of latency, and may be important in understanding CMV’s impact on immunity, inflammation and health over the lifespan.

## Results

### Neither dose nor age at infection influence viral loads in major sites of early infection

In order to investigate the influence of route and age on the course of mCMV infection from the acute to latent phase of infection, 4- (prepubescent) or 12-week (young postpubescent)-old male mice were infected with mCMV-Smith. Intraperitoneal (i.p.) infections were performed using 10^5^ p.f.u. while for i.n. infections we were limited to 10^4^ p.f.u. in 50 uL sterile saline. For the course of the experiment, we have used a combined longitudinal-cross-sectional design, with blood collected longitudinally, and the analysis in other organs being done in a cross-sectional manner (**Fig. 1A**). On d3 of infection, qPCR was utilized to measure the viral genomic load in various organs. We found variable, but not statistically significantly different, viral loads between different ages and routes of infection across different organs (**Fig. 1B**), allowing us to study differences in the host response in the face of similar virus spread and load in all groups of mice.

**Figure 1.**
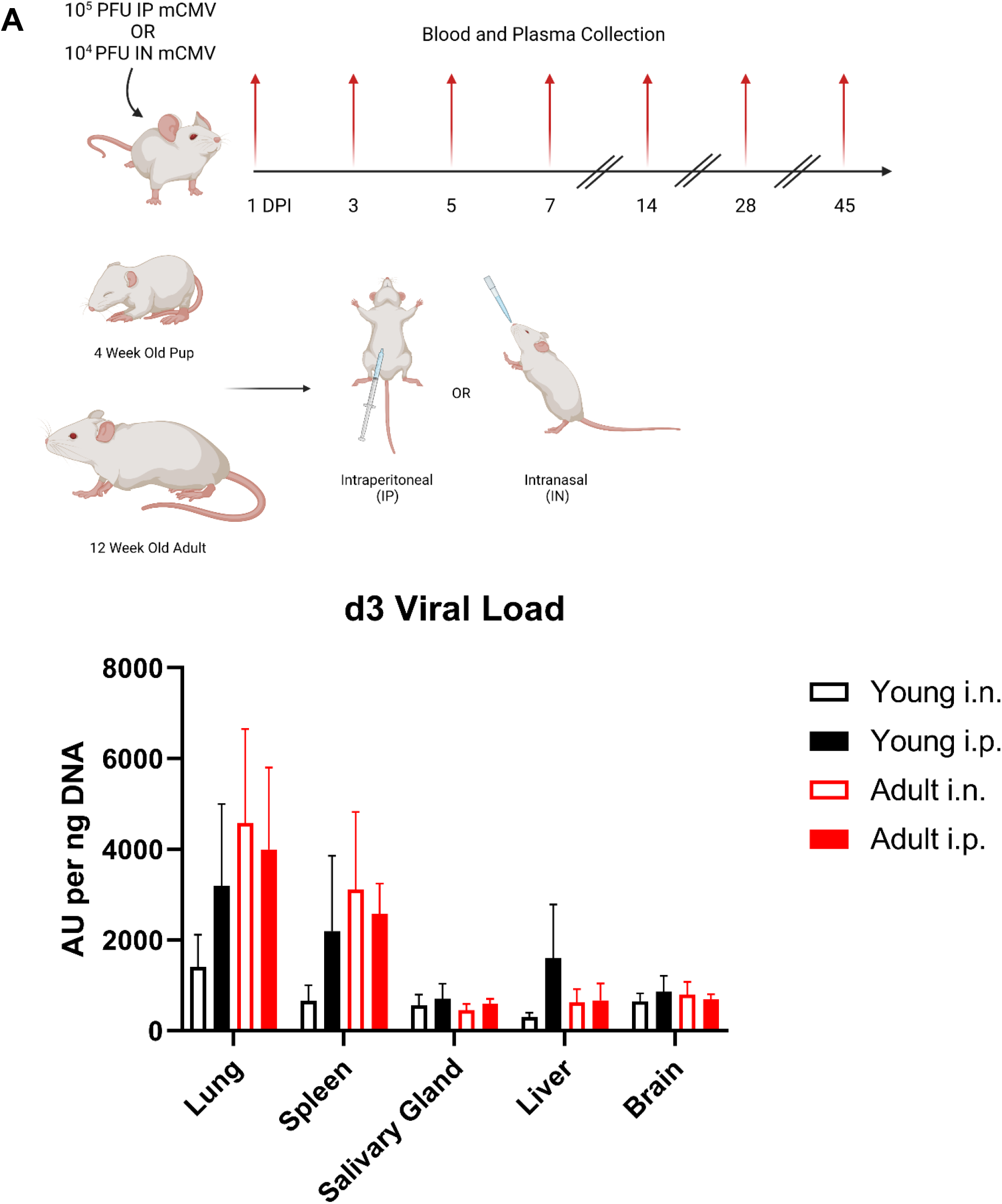
Neither dose nor age at infection influence viral loads in major sites of early infection.

### Both the route of, and age at infection affect activation and maturation of NK cells

Because of the known role of NK (CD3^-^NK1.1^+^) cells in controlling early mCMV infection^19^, we used multiparameter flow cytometry (FCM) to track the phenotype and kinetics of NK cell maturation and activation. We found that in all infection models the total number of NK cells in circulation rapidly contracted between d0 and d1, and then quickly expanded to maximal numbers on d7. While the numbers of NK cells in circulation in each model converge from d14 on, we found that the 4-week i.p. infected group expanded to a greater extent compared to other groups. This expansion in 4-wk i.p. infected mice outpaced the expansion in the other three groups on d3 and d5, and then receded to became similar to other groups by d7 (**Fig. 2A, right panel**).

**Figure 2.**
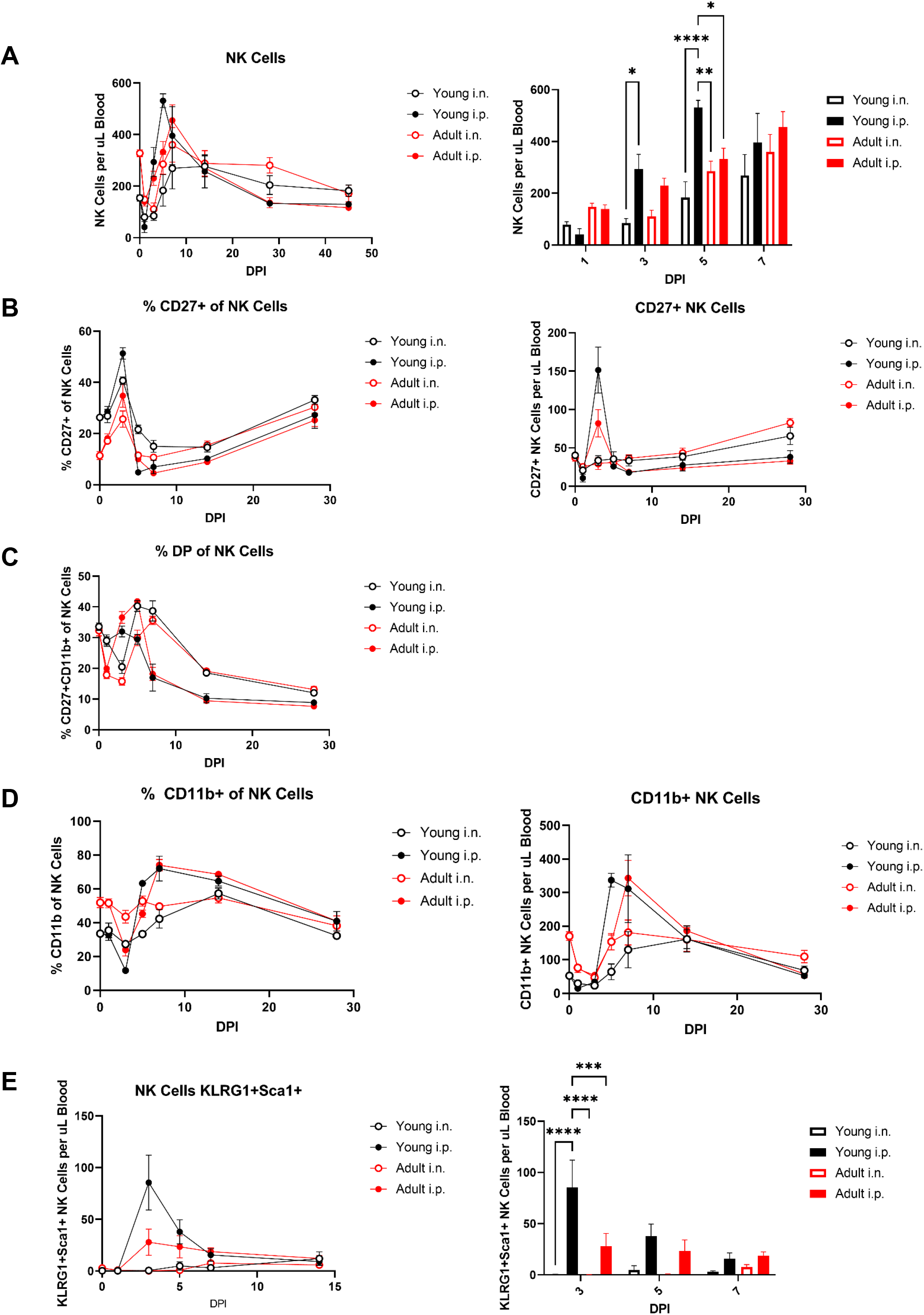
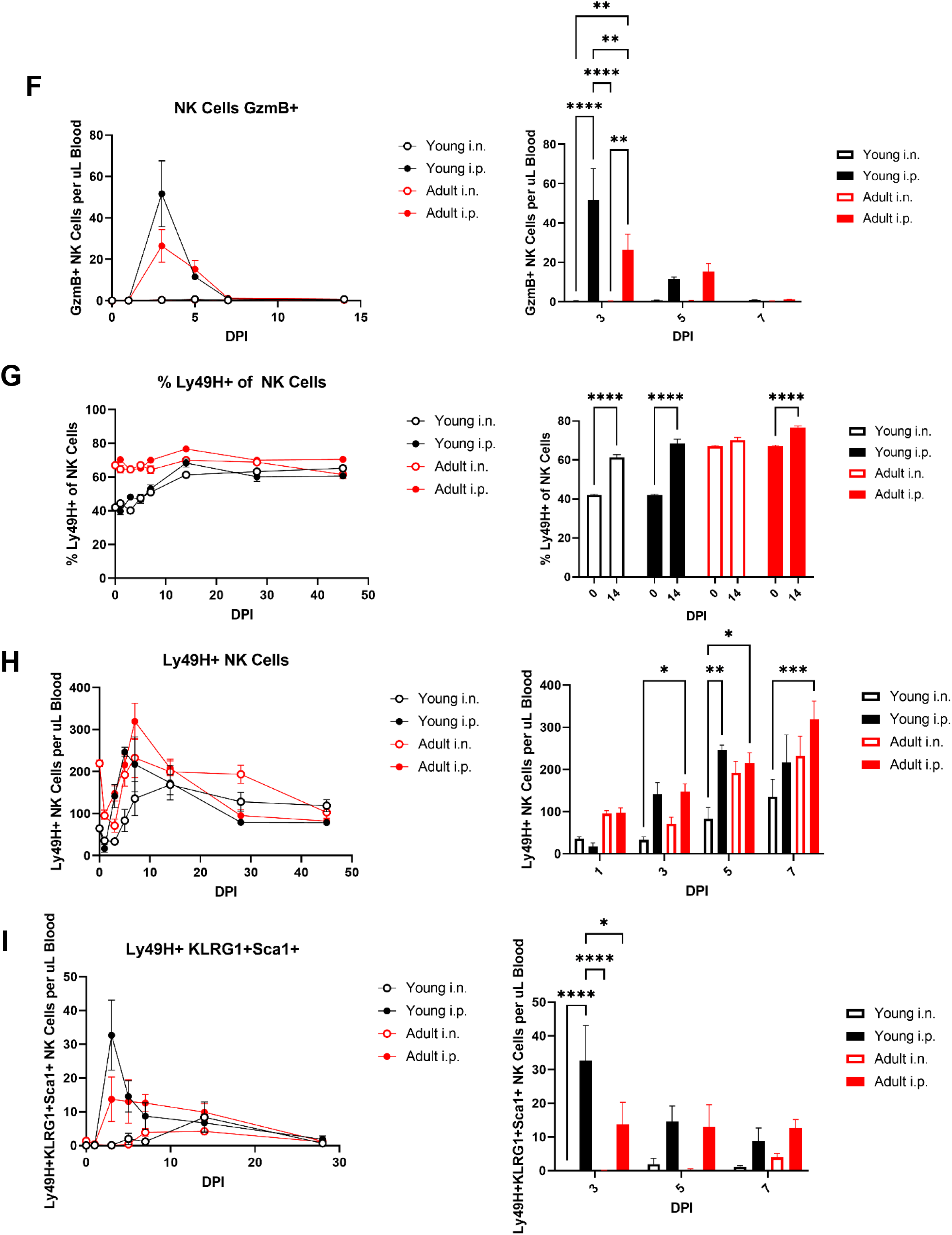
Both the route of, and age at infection affect activation and maturation of NK cells.

In both mice and man, the differential expression of CD27 and CD11b are known to subdivide NK cells into a four-stage developmental program from least mature CD11b^-^CD27^-^, via the CD11b^-^CD27^+^ and CD11b^+^CD27^+^ stages to the most mature CD11b^+^CD27^-20^. Evidence suggests that this progressive maturation is associated with the increased accrual of effector function. We found that prior to infection, 4-week old mice proportionally contained more immature CD11b^-^CD27^+^ NK cells (**Fig. 2B),** albeit these differences were not present when absolute cell counts were considered. By contrast, 12-week old mice contained both proportionally and absolutely larger population of mature CD11b^+^CD27^-^ NK cells (**Fig. 2D**). The NK cell compartment was modulated heavily by CMV in prepubescent mice, initially likely via maturation – a change of phenotype from less to more mature cells, as signified by the increased frequencies of double-positive cells soon after infection (d3) (**Fig. 2C**) and then their reduction by day 5-7 with a concomitant increase in single-positive CD11b^+^CD27^-^ mature NK cell frequency (**Fig. 2D**). Between days 5 and 7, there was an increase in frequencies of the least mature CD27+CD11b-cells, possibly due to their proliferation. Overall profiles of these changes were similar in postpubescent, 12-week old mice, but were somewhat less pronounced than those found in younger mice (**Fig. 2 A,B**). Towards the end of the first week, we found that NK maturation profiles segregated preferentially with the route of infection, with the largest proportion of most mature cells seen in i.p. infected mice, that were significantly more pronounced compared to i.n. infected mice regardless of age. At later times post infection (d28 and 45), all groups exhibited overlapping NK phenotype profiles and numbers in peripheral blood.

To further examine the activation status of NK cells in the blood during infection, we used Killer cell lectin-like receptor subfamily G member 1 (KLRG1) and Stem cells antigen-1 (Sca1). Approximately 30-40% of resting NK cells express KLRG1, while the expression of Sca1 is negligible. These two surface molecules are representative of early, non-specific NK cell activation^21^. While both KLRG1 and Sca1 were uniformly upregulated during early mCMV infection, KLRG1 expression persisted later into infection while the expression of Sca1 decreased rapidly prior to the end of the first week of infection. While there are reports suggesting that Sca1 may exhibit inhibitory function, Sca1 has been identified as a marker of functionally active NK cells – a higher proportion Sca1^+^ NK cells produce IFN-γ than their Sca1^-^ counterparts^22–24^. Co-expression of these two molecules therefore functionally identifies activated NK cells more stringently than the expression of either of these molecules alone. We found that in the case of i.n. infection, the number of bloodborne KLRG1^+^Sca1^+^ NK cells was negligible regardless of age. Conversely, the number of KLRG1^+^Sca1^+^ NK cells rapidly increased in the blood during i.p. infection, peaking at d3 and declining to a very small, but measurable, persistent population through the remainder of the infection. This is especially true in i.p. infected 4-week old mice, where the number of KLRG1^+^Sca1^+^ NK cells was ~3x higher that in 12-week old mice at the peak on d3 (**Fig. 2E**). A similar trend was seen in the expression of Granzyme B (GzmB), a cytotoxic serine protease-robust early production of intracellular GzmB, peaking on d3 and rapidly declining to baseline in i.p. infection, while i.n. infection did not yield a significant GzmB^+^ NK cell population in the blood. This was likely a consequence of regional immunity, whereby i.n. infection would likely elicit this population in the lungs and the salivary gland, an issue requiring further examination. Here too, i.p. infected 4-week old prepubescent animals on d3 possessed significantly higher (~2x) numbers of GzmB^+^ NK cells compared to i.p. infected 12-week old animals (**Fig. 2F**).

NK cells express an array of receptors that recognize major histocompatibility complex class 1 (MHC-I), that, among other functions, play a role in controlling viral infection. In mice, the Ly49 family of genes encodes such NK cell receptors. The activating Ly49H receptor is expressed on NK cells in genetically mCMV-resistant, but not susceptible, mouse strains (i.e., expressed in C57BL/6, but not BALB/c), and it has been shown to confer immune protection against mCMV infection, by binding to the virally encoded m157 protein to trigger NK cell cytotoxicity, cytokine and chemokine secretion^25^. We found that prior to infection ~65% of the total 12-week old mouse NK cell population expressed the Ly49H receptor, only ~40% of 4-week old NK cells were Ly49H^+^ (**Figure 2E**), that numerically translated to threefold higher absolute numbers of these cells (**Fig. 2F**). Numbers of Ly49H+ cells in the blood climbed quickly in the i.p. infection, peaking by d5-7. By contrast, in the i.n. infection the blood levels were initially barely detectable but caught up with the numbers seen in the i.p. infection by day 7 in both pre- and post-pubescent mice. Over the course of infection, however, the representation and numbers of Ly49^+^ NK cells increased in 4-week old mice much more than in 12-week old mice, making their frequencies and numbers largely comparable on or after d7 of infection for i.p. and d14 for i.n. infected animals. On d45 of infection, the proportion of Ly49^+^ NK cells out of the total NK cell population were indistinguishable regardless of age or route of infection (**Fig. 2E**).

When Ly49^+^ NK cells were examined for the expression of KLRG1 and Sca1, we found that on d3 of infection 4-week old i.p. infected animals had accumulated significantly more KLRG1^+^Sca1^+^ Ly49H^+^ NK cells than their 12-week old i.p. infected counterparts. At this timepoint, the number of these cells peaked, and by d5 of infection they declined to reach parity with their adult counterparts (**Fig. 2G**). Intranasally-infected animals did not exhibit an early accumulation of differentiated Ly49H+ KLRG-1+ Sca+ cells, however, these cells accumulated over time and by d14 and thereafter were represented at similar levels relative to the i.p. – infected mice in the blood.

These results indicate that the overall changes in activation and differentiation of Ly49H^+^ NK cells closely mirror that of the overall NK cell population during mCMV infection. Younger, prepubescent mice exhibited the strongest NK activation in the blood in response to i.p. infection, however, even the prepubescent 4-wk old i.n. infected mice accumulated the differentiated NK subtypes in the blood, albeit with somewhat delayed kinetics. The one exception were the NK cells expressing granzyme B, which did not appear at all in detectable numbers following i.n. infection in the blood. We hypothesize that in the i.n. infection, this cell type would be confined to the lungs and the salivary glands, as the site of initial encounter with the virus. Experiments to test this hypothesis are in progress.

### Intraperitoneal infection induces greater antigen-specific CD8 T cell and anti-mCMV IgG antibody responses

Next, we examined activation of the adaptive immune response as a function of different routes of mCMV infection at different ages. mCMV is known to elicit an antibody response, and the efficacy of this humoral response is well supported by experiments in which transferred serum from infected animals decreases the susceptibility of naïve mice to mCMV infection. During mCMV infection, levels of IgG have been demonstrated to rise during active infection, followed by stabilization once the active infection is resolved and the virus becomes latent^26,27^. Due to this, IgG antibody is commonly used to evaluate infection history in humans. We found the kinetics of the anti-mCMV IgG response in the serum to be initially similar across each age group and infection route. However, with time, and contrary to the NK cell responses, the differences between i.p. and i.n. infection became progressively larger, and with the older age correlating to higher Ab responses. Specifically, by d45 12-week old i.p. infected animals progressively accumulated a significantly larger amount of anti-mCMV IgG as compared to their 4-week old i.p. infected counterparts, whereas the responses in both i.n. infected groups seemed to have reached a plateau and did not further increase after day 28 (**Fig. 3A**).

**Figure 3.**
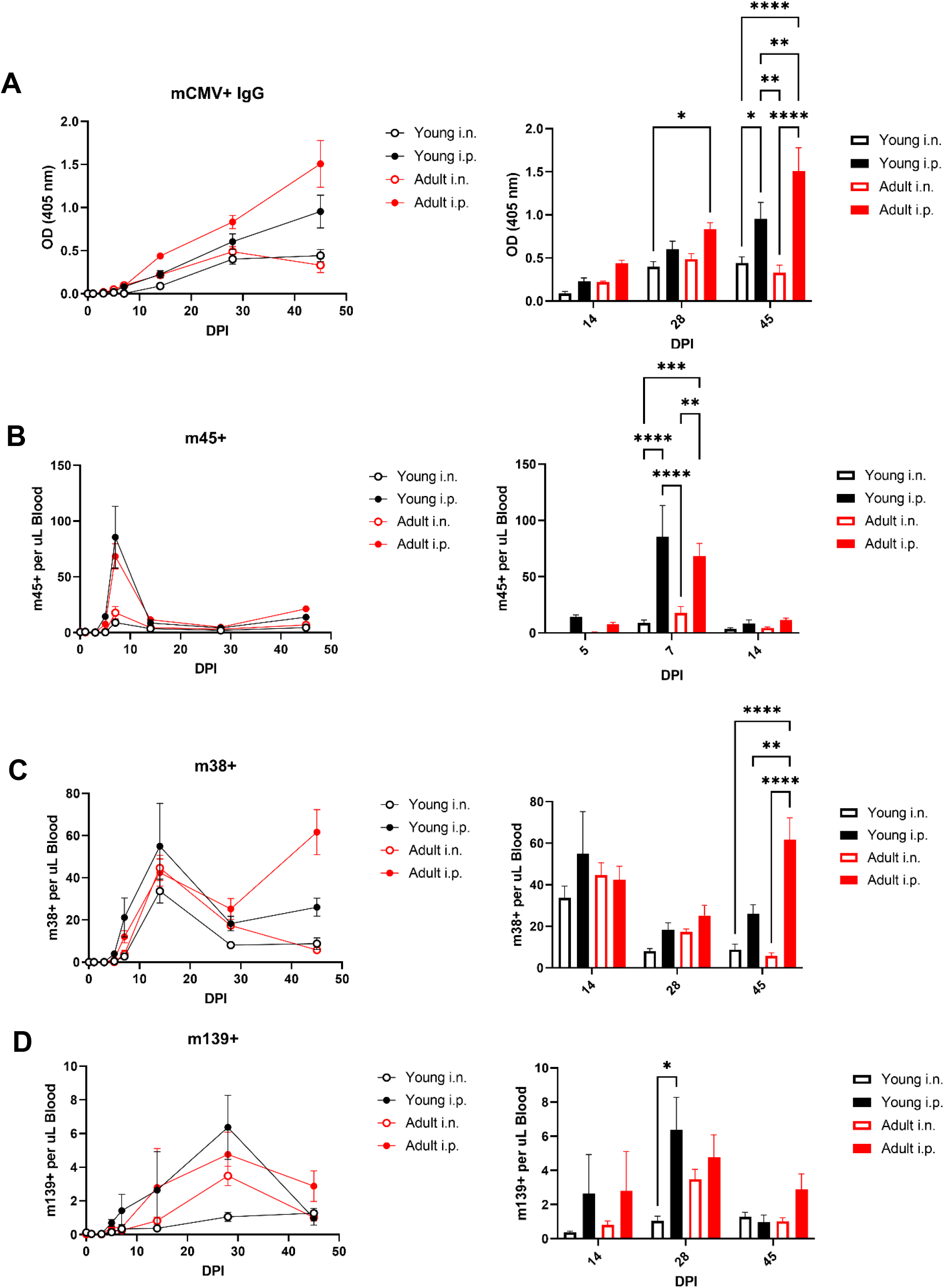
Intraperitoneal infection induces greater antigen-specific CD8 T cell and anti-mCMV IgG antibody responses.

One of the best-studied arms of adaptive immunity to CMV is the CD8^+^ T cell response. While CMV viruses encode several hundred proteins, exhaustive research in mice and humans has demonstrated that the CD8^+^ immune response targets vary as a function of the phases of infection, with certain epitopes eliciting strong responses in the acute phase, while those immunodominant in the latency/reactivation phases typically lead to memory inflation (“inflationary” epitopes)^28–30^. Thus, in B6 mice, the immunodominant protein of the acute anti-mCMV response is M45, a viral inhibitor of NF-kB^31,32^. We found that the expansion of M45-tet^+^ CD8^+^ T cells peaked in each group at d7, followed by a rapid contraction to a small but measurable population to d45. In both 4- and 12-week old mice, i.p. infected animals demonstrated a greater expansion of M45-tet^+^ CD8^+^ T cells in the blood, and this expansion was similar between both age groups (**Fig. 3B**).

Two of the hallmark inflationary CD8^+^ T cell responses in B6 mice are directed against the mCMV encoded proteins m38 (a mitochondrion-localized inhibitor of apoptosis) and m139 (an inhibitor of transcription, and production of type-I interferons) ^33–35^. We found that m38-tet^+^ CD8^+^ T cells were measurable by d7 of infection, increased in all groups by d14, then contracted in the two i.n. groups on d28 and 45, but not in i.p. groups, where after partial contraction by d28, they then increased significantly, with the 12-wk i.p. group showing the most significant inflation of all (**Fig. 3C**). m139-specific cells increased in all groups slowly to d28, after which point all groups showed contraction at d45. At that point, 12-wk-old i.p. infected mice showed a trend towards, but no statistically significant, increase in this response (**Fig. 3D**). These data suggest that the extent of systemic, bloodborne adaptive immune response varied in kinetics across age groups and infection routes depending on the type of the immune response, with markedly increased antibody and T cell responses in i.p. infected postpubescent animals.

### Overall changes in immune responsiveness to mCMV as a function of route and age at infection

**Figure 4** summarizes our findings on the measures of systemic innate and adaptive immunity against mCMV as a function of route and age at infection. We conclude that younger mice tended to exhibit increased NK cell numbers and differentiation following i.p. infection, whereas older i.p. infected mice exhibited stronger and prolonged, particularly inflationary, adaptive immune activation. Implications of these differences to lifelong immune homeostasis and immunity against CMV are discussed below.

**Figure 4.**
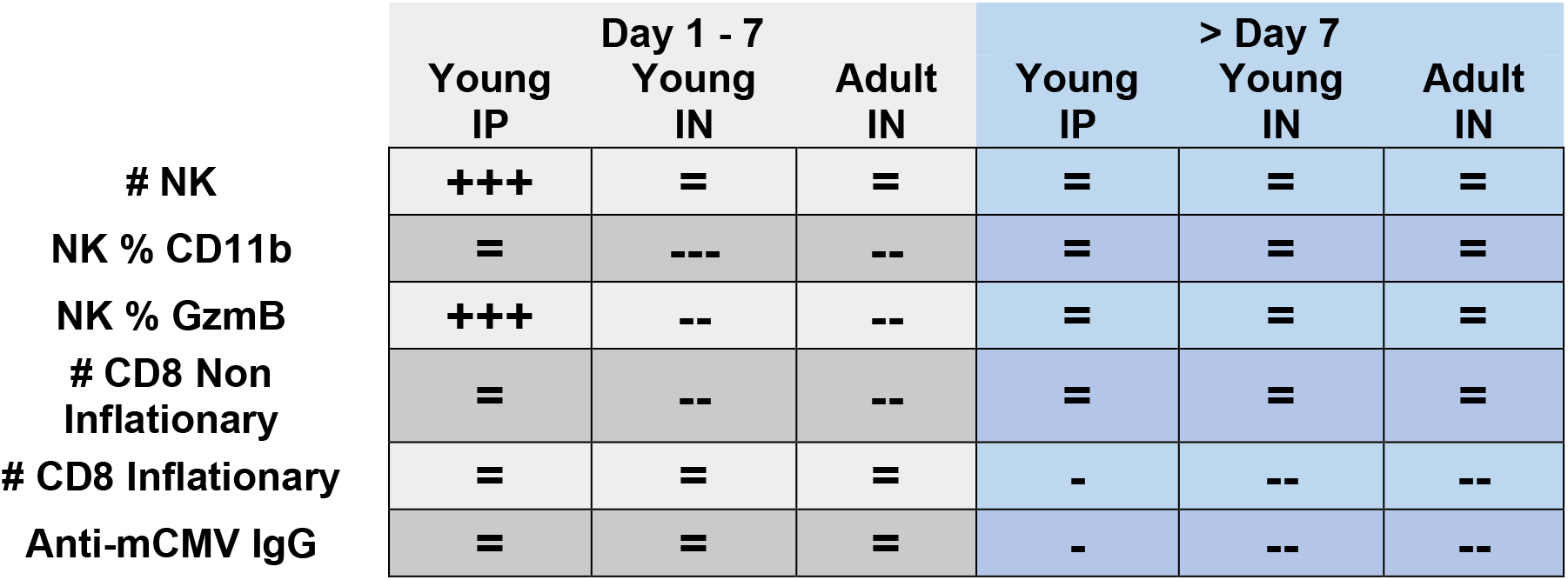
immune response to mCMV in the blood differs between infection routes and age groups in the deployment of cellular immunity. Comparison between age groups and routes of infection versus that measured in adult, i.p. infected animals. (=) no significant difference; (-) * lesser; (--) ** lesser; (---) *** lesser; (+) * greater; (++) ** greater; (+++) *** greater

## Discussion

Although the i.p. route of infection in 8-12-week old animals has been the most popular approach to the study of the biology of mCMV infection, it’s non-physiological characteristics make it imperative to assess whether other models may better approximate human infection. Our comparative analysis of both i.p. and i.n. routes of infection in 4-week (prepubescent) and 12-week (young postpubescent adult) old mice have revealed differences in immune control of mCMV as a function of the route and age at infection. While we ^36^ and others ^37^ have shown that mCMV infection results in local virus control via tissue-resident memory cells (Trm), it is critical for the organism to guard against systemic virus reactivation by maintaining a large pool of circulating T effector memory (Tem) and, in humans also Tem cells reexpressing CD45RA (Temra) cells in the blood.

The i.p. infection would be expected to result in a broader dissemination with faster and larger systemic responses, and that may not be representative of human infection, as it could distort many subsequent parameters of host:virus interaction, including the size of the cell pool for latency and reactivation, the extent and breadth of memory inflation and even the potential adverse health effects of CMV. Indeed, we found that the immune response to mCMV in the blood differs between infection routes and age groups in the deployment of different components of immunity (**Fig. 4A**), including differences in the NK, antibody and T cell responses.

Specifically, while the overall numbers of NK cells expanded until d7 and then receded in all infection groups, we found that less mature cells dominated in the i.n. infection. Age played a role in the NK activation as well, to the point that i.p.-infected 4-week old mice showed the greatest expansion and highest differentiation of NK cells, as judged by the expression of KLRG-1, Sca1 and granzyme B, all significantly higher compared to their i.p. infected 12-week old counterparts. Similarly, mCMV-specific Ly49H^+^ NK cells, that were represented at a lower frequency in prepubescent animals, accumulated quickly in the blood to dwarf those found in adult counterparts. Of interest, in i.n. infected mice, systemic NK responses were more visible at later times post infection, particularly around days 7 and later, typically catching up with those seen in i.p. infected mice, suggesting spread of the immune responses from the nasal/olphactory/respiratory immunity to systemic immunity.

These findings stand in contrast to lower, and with time less progressive, adaptive humoral and cellular immune responses, most notably the antibody and inflationary epitope CD8 T cell responses. These two types of responses were both progressive and by far most prominent on d45 in the i.p.-infected postpubescent young adult animals and were low in the i.n. infection groups. Experiments are in progress to track these parameters over time and in the presence or the absence of deliberate reactivation over the lifetime.

In humans, the consequences of CMV carriage in an immunocompetent host has been debated vigorously. In some studies, associations between CMV with all-cause death, cardiovascular diseases, diabetes, and other diseases of aging has been found, while in others, no such association was found^38^. Likewise, in mice, several studies have shown that mCMV may, albeit subtly, perturb various aspects of immune function in old age^39^, while other studies have shown benefit from CMV-carriage to the defense against other infections^40,41^. We hope that our present studies may provide initial impetus to define the optimal model of mCMV infection to ensure physiological applicability to hCMV infection. This should allow the field to conclusively reassess the possible link between the phenotypes found in causal experiments in mice to those in humans in the context of CMV, inflammation, healthspan and aging.

## Materials and Methods

### Ethics statement

Mouse studies were carried out in strict accordance with the recommendations in the Guide for the Care and Use of Laboratory Animals of the National Institutes of Health. Protocols were approved by the Institutional Animal Care and Use Committee at the University of Arizona (IACUC protocol 08-102, PHS Assurance No. A3248-01).

### Mice and mCMV infection

Adult (11 weeks old) and neonatal (3 weeks old) male C57/BL6 mice were obtained from Jackson Laboratories (Bar Harbor, ME). Mice were allowed to rest for 1 week following transfers between facilities. At 4 or 12 weeks of age, mice were infected with 10^4^ or 10^5^ p.f.u. of mCMV intraperitoneally or intranasally (Smith strain, obtained from M. Jarvis and J. Nelson, Oregon Health and Science University, Portland, OR). All mice were maintained in specific pathogen-free animal facilities at the University of Arizona (Tucson, AZ).

### Tissue and lymphocyte collection

Blood was collected into heparin-coated tubes and subjected to hypotonic lysis. 10 minutes prior to tissue collection, 100 μg CD45.2 antibody was administered i.p. in 100 μL in USP saline. Mice were euthanized using an overdose of isofluorane to maintain circulatory system integrity. Each tissue was harvested in RPMI supplemented with 5% FBS and finely minced. Tissue resident lymphocytes from the spleen (1), lungs (2), liver (3), and salivary gland (4) were isolated from their respective tissues as follows: (1) spleens were washed in RMPI with no FBS and subjected to Accutase treatment for 30 minutes at 37°C. Following digest, the mixture was filtered through a 40 μm mesh strainer and washed with 5% FBS RPMI. (2) Lungs were digested in 10 mLs of 5% FBS RPMI supplemented with .5 mg/mL collagenase type 1, .02 mg/mL DNase I, and 50 μL/mL elastase. After 30 minutes of incubation at 37°C, the digestion mixture was pushed through a 40 μm mesh strainer. Lymphocytes were isolated using Percoll density separation (30% and 70%) and washed. (3) Livers were pushed through a 40 μm mesh strainer and washed with 5% FBS RPMI. Lymphocyes were isolated using Percoll density separation (30% and 70%) and washed. (4) Salivary glands were subjected to digest in 8% FBS RPMI supplemented with 1 mg/mL collagenase IV, 5 mM CaCl_2_, 50 μg DNase I. After 30 minutes of digest at room temperature with continuous agitation, the digestion mixture was pushed through a 40 μm mesh strainer, washed with 5% FBS RPMI, and separated using a 30%/70% Percoll gradient.

### Flow cytometry and intracellular staining

Cells isolated from the blood and tissues were stained with surface antibodies and then fixed and permeabilized using the FoxP3 Fix/Perm kit (eBioscience, San Diego, CA). The following antibodies were used anti: CD3(145-2C11), CD4(GK1.5), CD8(53-6.7), CD44(IM7), CD62L(MEL-14), Ly6C(HK1.4), NK1.1(PK136), CD11b(M1/70), CD27(LG.3A10), CD28(37.51), KLRG1(2F1/KLRG1), Sca-1(D7), Ly49H(3D10), CD45.2(A20), Granzyme B (Biolegend, San Diego, CA), M45-tetramer, M139-tetramer, and M38-tetramer. Data acquisition was performed on a Cytek Aurora flow cytometer (Cytek Biosciences) and analyzed using FlowJo software (Tree Star, Ashland, OR). Gating was informed using fluorescence minus one (FMO) controls.

### Quantitative PCR of viral genome load in tissues

Spleen, liver, salivary gland, and liver were harvested from mCMV infected and uninfected age matched controls. Each tissue was collected into a microcentrifuge tube containing 1 mL of Qiazol (Qiagen) and autoclaved glass beads. Each sample was homogenized by bead beating with two 2-minute cycles. DNA was extracted per the Qiazol manufacturer’s protocol. qPCR was performed using PowerUP SYBR Green Master Mix (Applied Biosciences) on a Step One real-time PCR system (Applied Biosciences) using the following cycle protocol: an initial step at 2 min 50°C followed by 95° for 10 min, followed by 40 cycles of 95° for 15 sec, 60° for 1 min. Recombinant plasmids containing IE1 and C57/BL6 β-actin were used as template to establish standard curves for quantification. The primer sequences were as follows: IE1-fw (5’- CCC TCT CCT AAC TCT CCC TTT-3’) and IEI-rv (5’-TGG TGC TCT TTT CCC GTG-3’), β-actin-fw (5’-AGC TCA TTG TAG AAG GTG TGG-3’) and β-actin-rv (5’-GGT GGG AAT GGG TCA GAA G-3’). Cycle 32 was set as a negative cut-off based on uninfected controls. Primer sets and plasmid used to generate the standard curve were gifted by Wayne Yokoyama, MD, Washington University in St. Louis.

## Notes

Supported by USPHS award AG020719 and the Bowman Endowment in Medical Sciences to J.N-Z.

### Competing Interest Statement

The authors have declared no competing interest.

